# Baseline corticosterone does not reflect iridescent plumage traits in female tree swallows

**DOI:** 10.1101/452029

**Authors:** Christopher M. Harris, Oliver P. Love, Stéphanie M. Doucet, Pierre-Paul Bitton

## Abstract

The production of high quality secondary sexual traits can be constrained by trade-offs in the allocation of energy and nutrients with other metabolic activities, and is mediated by physiological processes. In birds, the factors influencing male plumage quality have been well studied; however, factors affecting female plumage quality are poorly understood. Furthermore, it remains uncertain which physiological traits mediate the relationship between body condition and ornaments. In this three-year study of after-second-year female tree swallows (*Tachycineta bicolor*), we investigated (1) the relationship between baseline corticosterone near the end of the brood-rearing period (CORT_BR_) and feather colour characteristics (hue, saturation, brightness) the following year, and (2) the relationship between baseline corticosterone measured during incubation (CORT_I_) and brood rearing (CORT_BR_), and feather colour in the same year. To control for reproductive effort, we included reproductive parameters as covariates in all analyses. In this first study between CORT and the plumage colour characteristics of a species bearing iridescent feathers, we did not find any relationship between CORT_BR_ and the colour of subsequently-produced feathers, nor did we find any relationship between CORT_I_ and the colour of feathers displayed during that breeding season. If CORT levels at the end of breeding carry over to influence the immediately subsequent moult period as we expect, our results generally indicate that structural plumage quality may not be as sensitive to circulating CORT levels compared to carotenoid-based colouration. Future studies, particularly those employing experimental manipulations of CORT during moult in species with iridescent traits, are necessary to fully determine the role glucocorticoids play in mediating the quality of secondary sexual characteristics.

## 1. Introduction

The honest signaling function of secondary sexual ornaments has been well established in a variety of species. Studies have shown that individuals bearing quality ornaments are in better body condition (e.g., Kodric-Brown and Brown, 1984), defend better/larger territories (e.g., Hill, 1988; Keyser and Hill, 2000; Wolfenbarger, 1999), provide better parental care (e.g., Hill, 1991; Linville et al., 1998; Siefferman and Hill, 2003), and have greater reproductive success and potential fitness (e.g., Bitton et al., 2007; McGraw et al., 2001). Signal honesty is likely to be maintained because individuals in better condition pay relatively lower costs for the development of sexually selected traits (Grafen, 1990; Rubenstein and Hauber, 2008), and costs of production are constrained by trade-offs with other metabolic activities and mediated through physiological processes (Tibbetts, 2014). In birds, most of the evidence for condition-dependent signaling has been provided through the study of male plumage (Dale, 2006; Griffith and Pryke, 2006; Hill, 2006; Senar, 2006). In contrast, relatively little is known about the condition-dependence of plumage colouration and the factors that may influence the production of quality feathers in females. This is of particular interest since male mate choice has been demonstrated in a large number of species (Amundsen, 2000; Amundsen and Pärn, 2006; Clutton-Brock, 2007), especially in cases where males and females both possess elaborate ornaments, and because females of most avian species invest more than males in reproduction (Kokko and Jennions, 2012). As a result, the plumage quality of females should reflect past reproductive investment and/or the potential for future reproductive effort, thereby influencing their ability to attract high quality mates (Clutton-Brock, 2009; Doutrelant et al., 2012).

The condition-dependence of plumage quality arises because of trade-offs in the allocation of energy and nutrients between the production of feathers and other costly metabolic and physiological maintenance activities. Exactly how these trade-offs are controlled is still largely debated (Morehouse, 2014). Because they are involved in the regulation and balance of energetic demands, physiological traits are thought to mediate the expression of traits advertising quality, including plumage colouration in birds (Bortolotti et al., 2009; Kimball, 2006; McGlothlin et al., 2008). Glucocorticoids, such as corticosterone (CORT) in birds, demonstrate great potential for this role: they are involved in the maintenance of whole-organism energetic balance (Landys et al., 2006), and mediate other key life history trade-offs (Crespi et al., 2013). Indeed, exogenously administered CORT has been found to slow the rate of feather growth (Romero et al., 2005), and higher levels of circulating CORT during feather growth can result in poorer quality feathers in terms of colouration, barbule density, strength, and micro-structure (DesRochers et al., 2009; Lattin et al., 2011; Roulin et al., 2008). Furthermore, levels of CORT in feathers, a potentially integrated measure of baseline CORT, have been found to correlate with carotenoid-based plumage colour (Kennedy et al., 2013; Lendvai et al., 2013), and baseline and stress-induced plasma CORT levels have been linked to measures of reproductive investment (Bonier et al., 2009a; Bonier et al., 2009b; Breuner et al., 2008; Love et al., 2014). Since moult often occurs immediately following breeding, plumage quality is likely to be influenced by past reproductive investment and success (Norris et al., 2004); however, whether CORT can mediate the longer-term trade-off between reproductive investment and feather quality is largely unexplored (Tibbetts, 2014), and has never been assessed for iridescent structural colouration.

The iridescent plumage of tree swallows (*Tachycineta bicolor*) is coloured by a single layer of keratin overlaying multiple layers of melanosomes in the barbules, which produces colour through thin-film interference (Maia et al., 2009; Prum, 2006). The mechanisms promoting the condition-dependence of iridescent plumage remain poorly studied (Doucet and Meadows, 2009; Maia and Macedo, 2011), and nanostructural differences leading to among-individual variation in colouration has been almost completely overlooked (but see Doucet et al., 2006). In satin bowerbirds (*Ptilonorhynchus violaceus*), a species in which structural colours are produced by the same nanostructural mechanism, the hue of the feather is associated with the average thickness of the keratin layer (Doucet et al., 2006), which is influenced by the size, quantity, and homogeneous distribution of melanosomes during feather keratinization (Maia et al., 2012). Colour purity (saturation) should be influenced by the homogeneousness of the keratin layer, and brightness has been correlated with barbule density (Doucet, 2002; Maia and Macedo, 2011). Because higher levels of CORT during feather growth have been demonstrated to reduce barbule density, strength, and micro-structure (DesRochers et al., 2009; Lattin et al., 2011; Roulin et al., 2008), and because physiological and nutritional stressors are known to induce keratin deposition abnormalities (e.g., fault bars; Jovani and Blas, 2004), baseline CORT levels during moult should influence iridescent plumage colouration. However, no studies have examined the potential relationship between iridescent plumage quality and circulating CORT.

In this study, we investigated whether the quality of mantle plumage colouration in after-second-year (ASY) female tree swallows reflects past or current year baseline plasma CORT. Using data collected over three years, we specifically test (1) the relationship between circulating CORT near the end of the brood rearing period (CORT_BR_) and feather colour characteristics (hue, saturation, brightness) the following year, and (2) the relationship between circulating CORT measured during the incubation (CORT_I_) and brood rearing period (CORT_BR_), and feather colour in the same year (**Figure 1**). We predicted that individuals with higher baseline CORT_BR_ would subsequently produce lower quality plumage, and that CORT_I_ and CORT_BR_ levels would be negatively correlated with current-year plumage quality attributes.

**Figure 1.**
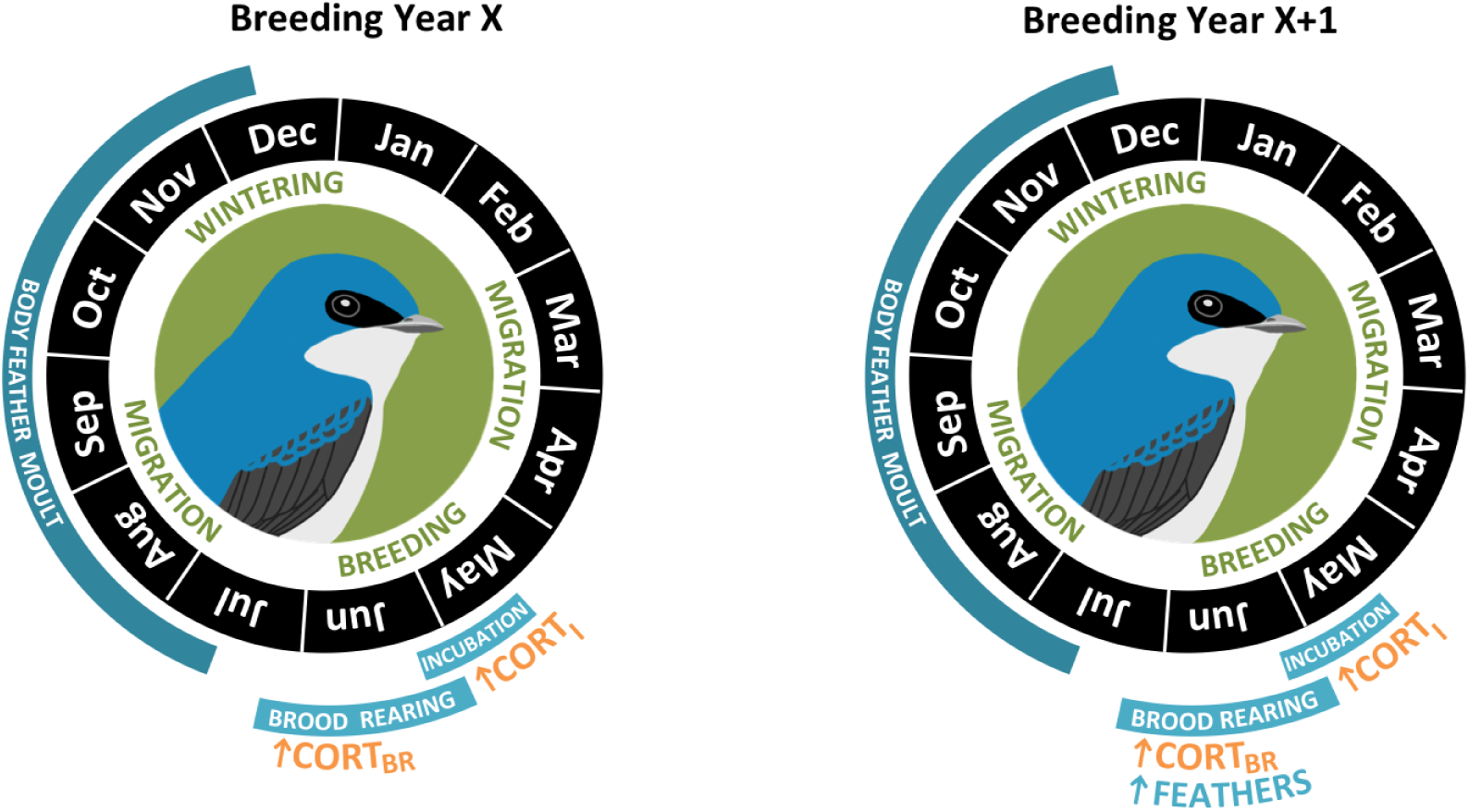
Sampling scheme for this study. Using a three year data set we tested the relationship between circulating CORT near the end of the brood rearing period (CORT_BR_; Year X) and feather colour characteristics (hue, saturation, brightness) the following year (Year X+1), and the relationship between circulating CORT measured during incubation (CORT_I_) and brood-rearing period (CORT_BR_) feather colour in the same year (e.g., Year X+l).

## 2. Methods

### 2.1 Species and study design

The tree swallow is a Neotropical migrant passerine that breeds across most of North America (Winkler et al., 2011). While second-year (SY) females possess dull grey plumage, males and ASY females bear iridescent upperparts which range in colour from green to blue among individuals (Hussell, 1983a). Delayed plumage maturation has been demonstrated to reduce conspecific aggression towards SY females during territorial intrusions, an adaptive trait in a species for which nest site competition is very high (Coady and Dawson, 2013). Even in iridescent adult females, reduced plumage brightness may reduce aggressive interactions (Berzins and Dawson, 2016). Male tree swallows with brighter plumage are older (Bitton and Dawson, 2008), and have greater extra-pair fertilization success (observational study in Bitton et al., 2007, and experimentally confirmed in Whittingham and Dunn, 2016). Bluer males (lower hue value) are older, have increased survival rates (Bitton and Dawson, 2008) and greater immune response (Beck et al., 2015) compared to greener males. In females, bluer feathers are associated with older females and greater fledging success (Bitton et al., 2008), and plumage brightness is positively correlated with total clutch egg mass (Bitton et al., 2008). Furthermore, plumage colouration seems to play an important role in intersexual interactions as there is assortative mating based on plumage brightness (Bitton et al., 2008), and individuals paired with greener mates (indicative of lower quality) feed their own nestlings at a greater rate (Dakin et al., 2016).

In general, tree swallows produce a single brood per season (Winkler et al., 2011). They are semi-colonial secondary cavity nesters that breed readily in artificial cavities and are highly philopatric (Winkler et al., 2004), which has made them an ideal study species for inter-annual investigations (Jones, 2003). In southern Ontario, breeding generally occurs from late April to mid-July and is usually immediately followed by moult (though moult can begin before nestlings fledge; see Hussell, 1983b). This complete prebasic moult begins with primary 1 and proceeds outward with body moult beginning with the back, breast, and belly regions when primary 2 to 4 are actively being replaced (Stutchbury and Rohwer, 1990). Moult continues through migration and is normally completed by mid-November (Stutchbury and Rohwer, 1990).

We studied two populations of tree swallows ^~^4 km apart in Haldimand County, Ontario: Ruthven Park National Historic Site 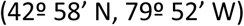 supports 140 nest boxes, and Taquanyah Conservation area 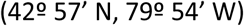 supports 35 nest boxes. Both areas are characterized by a matrix of riparian, agricultural, fallow field, and deciduous forest land use types. Tree swallows breeding at these locations have been comprehensively banded since 2010, and data for this study were collected during the breeding seasons (April to July) of 2011, 2012, and 2013. In each year, we monitored nest boxes daily to record nest-building activity, onset of laying, clutch size, and fledging success for all breeding pairs.

We captured ASY females inside their nest boxes during late incubation (10 days following clutch completion) and recorded body mass, wing length, and age, and banded unbanded birds with a standard Fish and Wildlife Survey aluminum leg band. Females were recaptured during the brood-rearing period (day 12 of nestling provisioning) and body mass was re-measured. This time period (approximately 7-9 days before fledging) coincides with maximum nestling mass (McCarty, 2001; Quinney et al., 1986), near maximum parental feeding effort (Leffelaar and Robertson, 1986), and the end of the safe period for handling nestlings without pre-fledging or affecting fledge date (Burtt, 1977). In 2011 and 2012, we also obtained a small blood sample from each female at each capture (Incubation - CORT_I_, Brood rearing - CORT_BR_) through puncture of the brachial vein (less than 10 % of total blood volume i.e., < 150 μl). We collected all blood samples between 0800 and 1200 hr to control for diel variation in CORT levels, and obtained all samples within two minutes of capture to ensure sampling of circulating baseline levels (Romero and Reed, 2005). Blood samples were stored on ice for up to five hours prior to centrifugation to separate plasma. Plasma samples were stored at −80° until assay. In 2012 and 2013, we also obtained five to six mantle feathers from each female during the incubation capture. Feathers were stored in opaque envelopes at room temperature until colour assessment.

### 2.2 Baseline corticosterone quantification

We determined plasma levels of total baseline CORT in non-extracted plasma using a Corticosterone Enzyme-linked-Immunosorbent Assay (ADI-900-097, EIA - Assay Designs, Michigan, USA; Madliger and Love, 2016a) with a 4-parameter logistic fit. This standard, commercially-available assay kit uses a sheep polyclonal antibody to corticosterone and 96-well plates coated with donkey anti-sheep IgG. Samples were run in triplicate at a total volume of 100 μl with 1:40 dilution and 1.5 % steroid displacement buffer. When measured concentrations fell below the detection limit of the assay (0.74 ng/ml), we assigned this detection limit to those samples (< 10 % of samples). The intra-assay variation was 8.0 % in 2011 and 10.3 % in 2012. Inter-assay variation was 13.3 % in 2011 and 6.0 % in 2012.

### 2.3 Feather characteristics

Five mantle feathers from each individual were overlaid as naturally found on a live bird and secured onto low-reflectance black velvet. Surface reflectance data was acquired using a USB2000 spectrophotometer and PX-2 Pulsed Xenon light source (Ocean Optics, Dunedin, Florida, USA) combined with a bifurcated fibre-optic probe. The probe was fitted with a rubber stopper at the tip to exclude ambient light, standardize the distance between the probe and the feathers, and allow each measurement to be taken at 90° from the surface (normal incidence). Five measurements were obtained from each sample; spectral data were recorded for wavelengths between 300 – 700 nm as the proportion of light reflected relative to the reflectance of a Spectralon pure white standard (Ocean Optics). Each spectral curve was smoothed using a locally-weighted scatterplot smoothing algorithm to prevent colour metrics from being influenced by effects of electrical noise. We subsequently averaged the five spectral curves, and calculated three colour metrics for the feathers of each individual. We calculated hue as the wavelength at maximum reflectance, mean brightness as the average reflectance over the 300 – 700 nm visual range, and chroma as the maximum reflectance minus the minimum reflectance divided by mean brightness (Andersson and Prager, 2006; Montgomerie, 2006). This measure of chroma generates increasing scores with increasing peak height while controlling for overall brightness (Andersson et al., 2002). Spectral curve processing and the extraction of colour metrics was conducted with the package ‘pavo’ (Maia et al., 2013) for the statistical language R (R Development Core Team, 2016). All data and scripts will be made available upon request.

### 2.4 Analyses

To investigate the relationship between CORT_BR_ levels and plumage characteristics following moult, we compared linear mixed models fit by restricted maximum likelihood. We used measures of CORT_BR_ and condition measured during the brood rearing period from year X (either 2011 or 2012) and measures of plumage colour from year X+1 (either 2012 or 2013; **Figure 1**). While CORT levels where not measured during moult, females in our population with higher baseline CORT during nestling provisioning returned the following year with higher baseline CORT during incubation (Madliger and Love, 2016a), indicating long-term repeatability of individual CORT levels. As a measure of body condition, we used body mass, because the residual of a linear regression of mass over wing length was not significant (year included as covariate: F_2,32_ = 1.06, p = 0.37; Schulte-Hostedde et al., 2005). To avoid pseudoreplication by including values for females that had been captured during more than one inter-annual breeding attempt, all models included female identity as the within-subject random factor. For each of the three plumage characteristics included as the dependent variable (hue, brightness, chroma), we generated a set of eight biologically relevant candidate models (and an additional intercept-only ninth model) which were compared using Akaike’s information criterion corrected for small sample size (AICc; Akaike, 1973; Burnham and Anderson, 2002; Hurvich and Tsai, 1989). All models except for the intercept-only model included CORT_BR_ and year of data collection. Seven of the models also included as independent factors body mass, laying date, and clutch size in all possible combinations (See Supplemental Material for the complete set of models). We did not include female age because the full study site was initiated the year before this study began, and we had incomplete age data for too many individuals. The best fitting models were considered equally plausible when the AICc value differed by no more than 2.00 (ΔAICc < 2.00) compared with the model with lowest value (Burnham and Anderson 2002). Variance explained by the best fitting models was calculated using marginal and conditional R^2^ (Nakagawa and Schielzeth 2013) using the ‘rsquared’ function in the R package ‘piecewiseSEM’ (Lefcheck, 2016).

To investigate the predictive value of plumage characteristics on CORT_I_ and CORT_BR_ levels, we compared linear models using measures of plumage characteristics, condition, and CORT collected within the same year (2012 only). For each of the two CORT measures we compared a set of candidate models that all included in various combinations at least one of the three plumage characteristics (hue, brightness, chroma). In half of the models, two measures of condition, laying date (Verhulst and Nilsson 2008) and body mass, were also included. The regression of mass over wing length was also not significant for this group of females (incubation mass: p = 0.08; brood rearing mass: p = 0.20). All sets of models included a global model and an intercept model (See Supplemental Material for the complete set of models), and the best fitting models were selected based on their AICc scores as above.

## 3. Results

When investigating the relationships between CORT_BR_ levels and plumage characteristics following moult, we used 37 observations obtained from 33 different females (four females captured in both years). To satisfy assumptions of normal distribution, we log-transformed CORT_BR_. We treated clutch size as an ordinal factor, including only nests that had between four and seven eggs to avoid strong imbalance in sample size between groups. Before comparing models, we ensured that they did not suffer from multicollinearity of independent factors through the calculation of variance inflation factors (VIFs). The set of models that best fit the data for plumage brightness and chroma had the intercept-only model indicating that none of the models fit the data well (**Table 1**). A single model fit the data for plumage hue, and included the predictors CORT, clutch size and mass (marginal R^2^ = 0.132, Conditional R^2^ = 0.132). However, only mass (estimate: −10.01; 95 % CI: −19.21 – −0.81) but neither CORT (estimate: −0.88; 95 % CI: −10.40 – 8.63) nor clutch size (estimate: −1.25; 95 % CI: −22.66 – 20.16), were significant predictors.

**Table 1.**
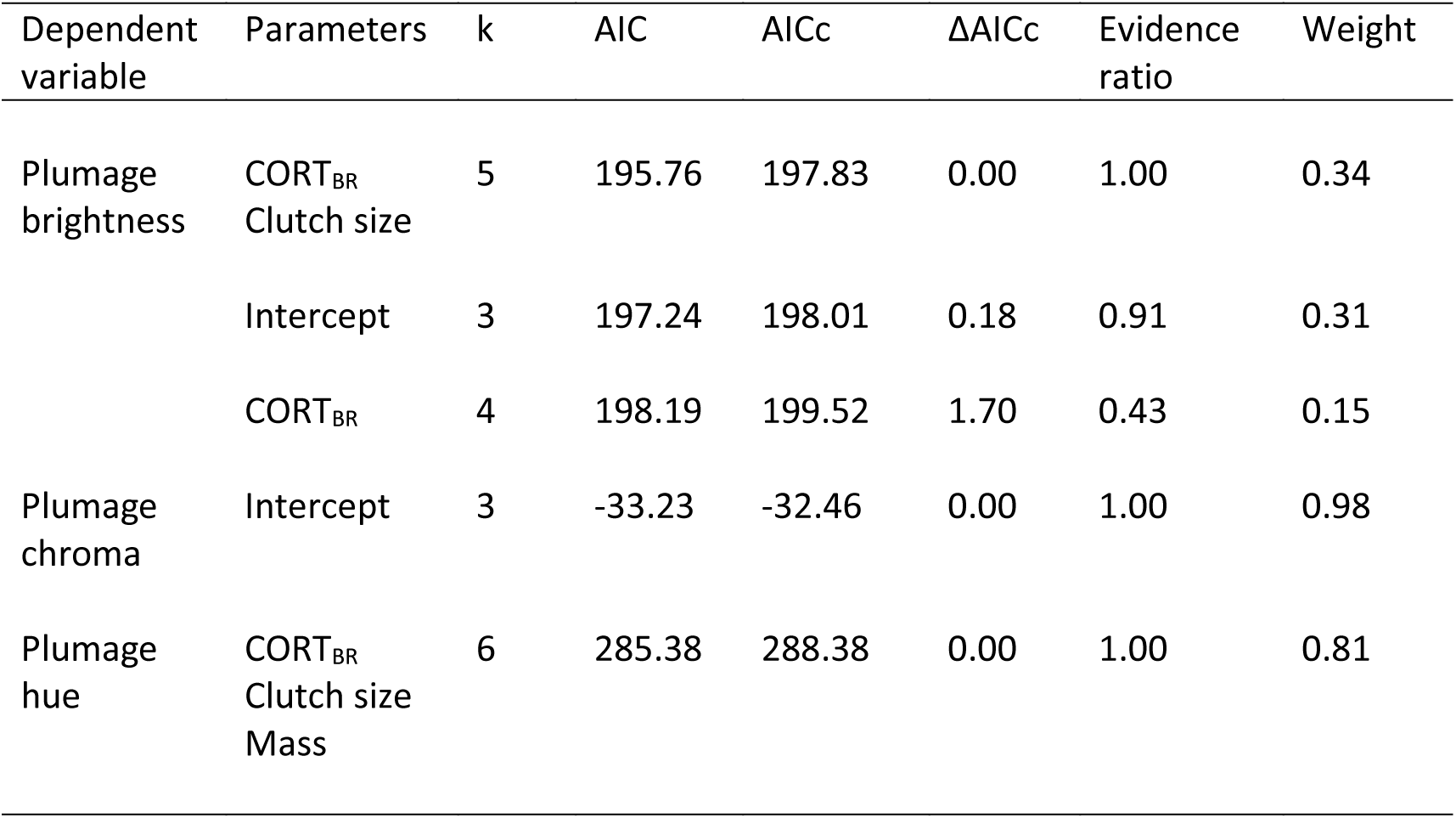
Summary of the linear mixed models that best fit the data when investigating the relationship between baseline corticosterone sampled during the brood-rearing period (CORT_BR_) in year X and plumage characteristics sampled in year X+1 (these feathers would have grown in shortly after brood rearing when corticosterone was sampled the previous year). All models also included the year of feather collection as covariate (not show for brevity). From all candidate models, only those with a ΔAICc < 2.0 are presented.

When investigating the relationships between our two corticosterone measures (CORT_I_ and CORT_BR_) and same year plumage characteristics, we used 32 observations, all from different females. CORT_I_ and CORT_BR_ were log transformed, only nests that included between four and seven eggs were used, and clutch size was treated as an ordinal factor. We also ensured that our models did not suffer from multicollinearity. The set of models that best fit the data for CORT_I_ included the intercept-only model indicating that none of our models fit the data well (**Table 2**). In contrast, two models fit the data for CORT_BR_. The first model included brightness, female body mass and clutch size as predictors (adjusted R^2^: 0.27); the second included brightness, hue, female body mass and clutch size (adjusted R^2^: 0.27; **Table 2**). We calculated the parameter estimates by weighting the parameter values and associated standard deviations of the two equally likely models by their respective model weight. This indicated that neither brightness (estimate: −0.065; 95 % CI: −0.139 – 0.008), hue (estimate: 0.005; 95 % CI: −0.005 – 0.015), nor body mass (estimate: −0.125; 95 % CI: −0.352 – 0.103), were significant predictors. In these models only clutch size (estimate: 0.96; 95 % CI: 0.41 – 1.51) was a significant predictor.

**Table 2.**
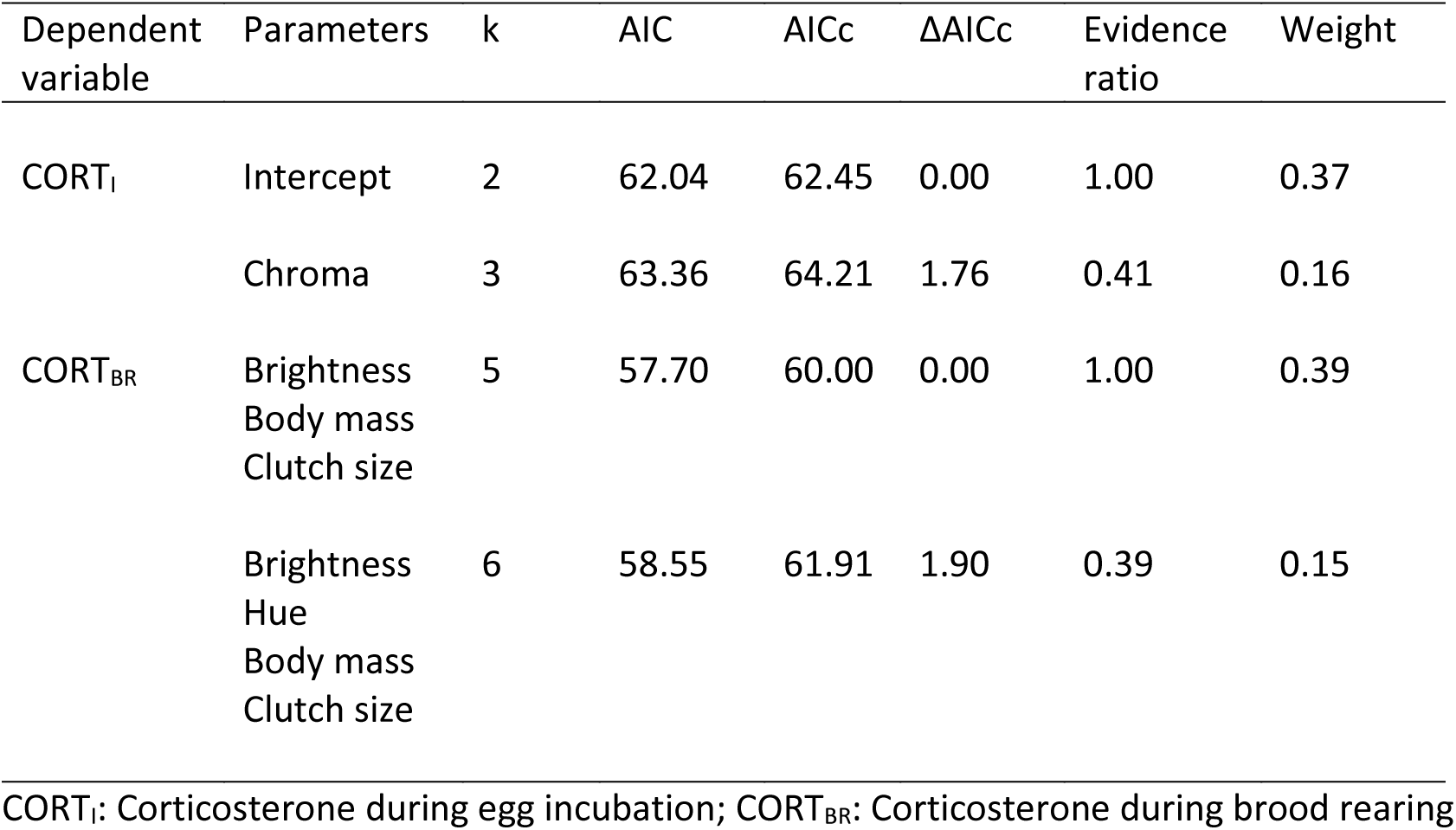
Summary of the linear mixed models that best fit the data when investigating the relationship between same-year corticosterone levels collected during the incubation period (CORT_I_) and brood-rearing period (CORT_BR_), and plumage characteristics. From all candidate models, only those with a ΔAICc < 2.0 are presented.

| Dependent variable | Parameters | k | AIC | AICc | $\Delta AICc$ | Evidence ratio | Weight |
| --- | --- | --- | --- | --- | --- | --- | --- |
| $CORT_I$ | Intercept | 2 | 62.04 | 62.45 | 0.00 | 1.00 | 0.37 |
|  | Chroma | 3 | 63.36 | 64.21 | 1.76 | 0.41 | 0.16 |
| $CORT_{BR}$ | Brightness<br>Body mass<br>Clutch size | 5 | 57.70 | 60.00 | 0.00 | 1.00 | 0.39 |
|  | Brightness<br>Hue<br>Body mass<br>Clutch size | 6 | 58.55 | 61.91 | 1.90 | 0.39 | 0.15 |

$CORT_I$ : Corticosterone during egg incubation; $CORT_{BR}$ : Corticosterone during brood rearing

## 4. Discussion

There is mounting evidence in some species that female plumage colouration is not a genetic carryover from male plumage colouration, that female plumage characteristics are associated with quality and reproductive success, and that males select females based on secondary sexual characteristics (Amundsen, 2000; Amundsen and Pärn, 2006). However, we know very little about which physiological and/or metabolic processes link the cost of reproduction to plumage quality in females (Moore et al., 2016). In this study, we used data collected over three years on breeding ASY female tree swallows to investigate the relationship between individual variation in baseline plasma corticosterone (CORT) and the characteristics of iridescent plumage. We predicted that females with high baseline CORT levels just prior to moulting would produce low quality plumage, and that females with high quality plumage would have low baseline CORT upon returning to the breeding site. In contrast to our predictions, plumage characteristics were not associated with previous or subsequent circulating CORT levels.

### 4.1 Baseline corticosterone as a predictor of plumage quality

Because moult occurs immediately following breeding in tree swallows (Winkler et al., 2011), the large investment in reproduction is expected to limit the energy and resources available for the production of high quality feathers (e.g., Hemborg and Lundberg, 1998; Norris et al., 2004). Indeed, observational and experimental studies have demonstrated the influence of breeding effort on structural plumage quality in other species. For example, male, but not female, eastern bluebirds (*Sialia sialis*) raising artificially enlarged broods produced lower quality non-iridescent plumage (Siefferman and Hill, 2005, 2008). Similarly, the plumage hue of male and female blue tits (*Cyanistes caeruleus*) shifted towards shorter (bluer) wavelengths, an indication of higher quality plumage, after fledging regular size broods, but not enlarged broods (Doutrelant et al., 2012). Because reproductive effort has also been associated with increased baseline CORT in several studies (Crespi et al., 2013), we predicted that females with higher levels of CORT immediately preceding feather moult (i.e., CORT_BR_) would produce lower quality plumage. Unlike these previous studies, our research failed to find any relationship between CORT_BR_ and subsequent plumage colouration. However, it is important to note that all studies presented above were experimental in design, whereas we did not manipulate the reproductive effort of the females included in our study. Ours is but a first step in identifying the signaling mechanisms that mediate plumage colour in female tree swallows, but experiments that either directly increase CORT levels during moult or manipulate reproductive effort could arrive at different conclusions. Other non-mutually exclusive reasons may explain our results.

Baseline CORT as assessed in this study is a temporally-specific measure which can vary based on a number of factors including social interactions (Creel, 2001), food availability (Jenni-Eiermann et al., 2008), and predation risk (Cockrem and Silverin, 2002). Because this measure of energetic stress is known to vary on a short temporal scale, it may not be an accurate reflection of the condition of the individual at the time of feather production. Although it was not possible to obtain CORT measures from moulting birds, and we compared plumage characteristics to CORT measurements obtained as close as possible to the production of the new feathers, a change in the phase of the annual cycle from breeding to migration could influence the levels of plasma CORT during the production of feathers (Romero, 2002). However, birds with higher CORT levels in one life history stage (e.g., breeding) could be expected to also have higher CORT levels during subsequent stages (e.g., moult). While baseline CORT levels generally show low repeatability between years, they can show moderate within-individual repeatability over shorter time scales (weeks to months) (Ouyang et al., 2011; Romero and Reed, 2008; Schoenemann and Bonier, 2018; Taff et al., 2018). Even over a longer time frame, in our population of tree swallows, females with higher baseline CORT during nestling provisioning return the following year with higher baseline CORT during incubation (Madliger and Love, 2016a). Therefore, we believe it is possible that females with elevated CORT at the end of the nestling provisioning stage could continue to experience elevated CORT during the moult period, which occurs immediately following the cessation of breeding (i.e., females that find one stage energetically-demanding may also face higher physiological stress in subsequent stages).

If the production of quality plumage is important for females, as would be implied by assortative mating found in this species (Bitton et al., 2008), female tree swallows could be down-regulating their CORT levels during feather replacement to avoid the negative effects on protein production and therefore feather quality. CORT levels in some species are known to be at their lowest towards the end of the breeding season and during moult (e.g. Done et al., 2011; Romero, 2002), and CORT-implanted European starlings (*Sturnus vulgaris*) have been shown to decrease feather growth leading to the production of feathers with quality independent of CORT levels (Romero et al., 2005). Indeed, CORT levels measured in feathers are not always associated with plasma CORT (Patterson et al., 2015), suggesting the existence of a mechanism that would control CORT levels in follicles. As a result, there may be inter-individual variation in the ability to down-regulate CORT production during moult (Romero, 2002) or in the release of CORT into the growing feather (Harris et al., 2016), which could lead to the lack of relationship between plasma CORT at nestling provisioning and feather quality indices.

While the feathers were produced soon after the collection of the CORT_BR_ sample, they were not collected for characterization until about nine months later, once the females had returned to the breeding grounds. During this time, feather characteristics may have changed. The brightness of plumage is known to be reduced through abrasion in carotenoid-based (Figuerola and Senar, 2005; McGraw and Hill, 2004) and non-iridescent structurally coloured feathers (Örnborg et al., 2002), and it is thought that wear may also be responsible for changes in hue and saturation of non-iridescent structural colouration (Örnborg et al., 2002). If changes in feather colour characteristics were non-random in such a way that higher quality feathers were more likely to be degraded, a relationship between CORT_BR_ and feather quality the subsequent year would be more difficult to detect. However, it may be more likely that individuals that produced high quality plumage can allocate more energy to feather maintenance, thus reducing degradation, which would increase the perceived effect of CORT_BR_ on feather quality.

Finally, our inability to accurately age females may have added confounding variation to our study. Since reproductive investment and reproductive success are influenced by age in tree swallows (Robertson and Rendell, 2001), and that these factors (including age) have been associated with plumage colouration (Bitton and Dawson, 2008; Bitton et al., 2008), it may be difficult to disentangle the relationships between reproduction-induced stress, age, and plumage attributes without specific age information. For instance, it is possible that younger females have more difficulty managing stress or energetic demand than older females (Wingfield and Sapolsky, 2003), which could lead to relationships between plumage traits and CORT in young females but not in older females. A longitudinal study, or experiment in which CORT levels are manipulated while blocking treatments on female age, could help resolve these questions.

### 4.2 Plumage characteristics representing current-year corticosterone

The role of plumage colouration in inter-and intra-sexual competition has been well documented (Griffith and Pryke, 2006; Senar, 2006). A common premise is that plumage characteristics are representative of an individual’s quality even if those feathers were produced months before the arrival to the breeding grounds. Since baseline CORT levels have been experimentally shown to mediate reproductive investment in birds (e.g., Bonier et al., 2011; Hennin et al., 2016; Love et al., 2014), we predicted a negative relationship between same year plumage colour attributes and CORT during reproduction. Our lack of significant results may not be surprising as a few studies have found a potential role for CORT as a mediator between condition and plumage colouration (e.g., Fairhurst et al., 2014; Grindstaff et al., 2012; Lendvai et al., 2013), while several have not (e.g., Jenkins et al., 2013; Merrill et al., 2014). Even among studies that have found relationships between CORT and secondary sexual characteristics, the direction of the relationship is not always as predicted (negative association between CORT levels and plumage quality), and may be context-dependent (Fairhurst et al., 2014; Lendvai et al., 2013). Furthermore, given the correlative nature of the current study it is possible that the individuals were not subjected to sufficient stress to lead to effects on ornament quality. Direct or indirect manipulation of circulating CORT right at the end of the breeding season or during moult, or handicapping the birds to increase their work load, would help determine its relationship with iridescent plumage quality.

Unlike the large majority of studies investigating relationships between CORT and plumage attributes, tree swallows display structurally-coloured plumage, not pigment-coloured plumage (see Grindstaff et al., 2012 for an example of non-iridescent structural study). Keratin and melanin, which produce the nanostructures responsible for the colours in tree swallow feathers, are metabolically produced *de novo* at the follicle during feather development (McGraw, 2006b), unlike carotenoids which must be acquired from the diet and used by other physiological processes (e.g., anti-oxidants, McGraw, 2006a). Therefore, keratin and melanin may not be influenced by metabolic processes in the same way as carotenoids (Fairhurst et al., 2015). Indeed, studies investigating the link between CORT and plumage attributes have more often found relationships in species bearing carotenoid-based plumage, not melanin-based plumage. In mallards, *Anas platyrhynchos*, feather CORT in ducklings was positively correlated with carotenoid-based signals in adults, but not correlated with melanin-based signals (Fairhurst et al., 2015). Similarly, in yellow warblers, *Setophaga petechia*, feather CORT negatively correlated with several measures of carotenoid-based plumage signals (hue, chroma), but not phaeomelanin-based plumage attributes (Grunst et al., 2015). Several other studies have failed to find relationships between CORT and melanin or structurally based plumage colouration (e.g., Jenkins et al., 2013, but see Grindstaff et al., 2012 and Henderson et al., 2013 for contrary examples). The potential for energetic and physiological costs to influence different aspects of plumage colouration likely depends on the underlying physical and pigmentary structures that are responsible for the colours, how their organization is controlled in the developing feathers, and feather maintenance behaviours.

It is possible that the level of reproductive demand and baseline CORT affects aspects of plumage colouration that we did not measure. Iridescent plumage, by definition, changes in colour with changes in angle between the viewer and the light source. Iridescent feathers could therefore be assessed for the range of angles over which colour is displayed (called *directionality* in Van Wijk et al., 2016b), and the hues at these angles. These metrics may reflect different feather properties than those obtained from taking measurements at normal incidence (hue, saturation, and brightness calculated at 90° from the feather surface), and *directionality* better predicts some aspect of male reproductive success in this species. Future studies on the proximate mechanism of iridescent plumage colouration should assess these changes in colour with viewing angle and better evaluate the functionality of iridescence itself (Meadows et al., 2011; Van Wijk et al., 2016a).

### 4.3 Conclusion

The role of variation in baseline CORT in mediating the relationship between condition and secondary sexual characteristics in birds is far from being well understood. A large number of factors can influence CORT levels at any time (Moore et al., 2016), and measures of stress are not always repeatable within individuals between seasons and years (Legagneux et al., 2013; Perez et al., 2016; Schoenemann and Bonier, 2018; Taff et al., 2018). In an attempt to best capture a temporally valid measure of stress, we collected blood samples to assess baseline plasma CORT during the incubation period and during the brood rearing period. This allowed us to compare the plumage characteristics to the most relevant levels of CORT: CORT_BR_ to plumage traits produced soon after leaving the nest, CORT_I_ to plumage colouration upon arrival at the breeding site. However, we did not find any relationship between CORT levels and plumage traits in female tree swallows. While multiple explanations can be invoked to rationalize our findings, the interpretation of our results is difficult because there is a lack of studies investigating the link between CORT and the production of ornaments in females, and a paucity of studies investigating the link between CORT and the production of structural colouration, both iridescent and non-iridescent. It is generally unknown how trade-offs between plumage traits and metabolism are mediated differently in males and females, and it is not clear what relationship we should expect between CORT and the expression of colours produced from nanostructures. This invites future studies on the topic, particularly ones that would manipulate CORT levels (or workload that would elevate baseline CORT; Madliger and Love, 2016b) in individuals.

## Acknowledgements

We thank Ruthven Park National Historic Site, the Grand River Conservation Authority, and Habitat Haldimand for access to field sites. We also thank three anonymous reviewers for helpful comments on the manuscript.

## Author contributions

K.S., C.L.M., and P-P.B. contributed equally to this paper. K.S., C.L.M., and P-P.B. conceptualized the study. K.S, C.L.M., and C.H. collected data. C.L.M., and P-P.B. analyzed the data. All authors contributed to writing the manuscript and to editing the final version.

## Declarations of interest – None

Permits – All procedures were approved by the University of Windsor’s Animal Care Committee (AUPP #10-10) and the Canadian Wildlife Service (Permit CA 0266).

